# Quality control of *Bradyrhizobium* inoculant strains: Detection of *nosZ* and correlation of symbiotic efficiency with soybean leaf chlorophyll levels

**DOI:** 10.1101/2023.10.26.564198

**Authors:** Damián Brignoli, Emilia Frickel-Critto, Tamara J. Sandobal, Rocío S. Balda, Cecilia B. Castells, Elías J. Mongiardini, Julieta Pérez-Giménez, Aníbal R. Lodeiro

**Affiliations:** Instituto de Biotecnología y Biología Molecular (IBBM)-Facultad de Ciencias Exactas, Universidad Nacional de La Plata y CCT-La Plata, CONICET, Calles 47 y 115 (1900) La Plata, Argentina; Cátedra de Genética-Facultad de CienciasAgrarias y Forestales, Universidad Nacional de La Plata, Calles 60 y 119 (1900) La Plata, Argentina; Laboratorio de Investigación y Desarrollo de Métodos Analíticos, Universidad Nacional de La Plata y CIC-PBA, Calles 47 y 115 (1900) La Plata, Argentina

**Keywords:** *Bradyrhizobium*, soybean, chlorophyllometer, nitrogen fixation, nitrous oxide

## Abstract

Greenhouse gas emissions, such as N_2_O from excessive N-fertilizer use, are of concern. Symbiotic N_2_-fixation by pulses as soybean might mitigate this issue, for which inoculants carrying locally adapted *Bradyrhizobium* strains are recommended. In the frame of this goal, enhancing the quality control of these inoculants is required on two key aspects: determining the presence of *nosZ*, for the strains being able to reduce N_2_O, and assessing N_2_-fixation potential. Yet, simple and non-destructive methods to assess N_2_-fixation are lacking. Here we aimed to leverage the correlation between N and chlorophyll levels by cultivating soybeans in vermiculite with N-free nutrient solution, inoculating them with various *Bradyrhizobium* field isolates, and subsequently measuring chlorophyll with a portable chlorophyllometer, correlating it to symbiotic parameters. We observed significant correlations between chlorophyll and shoot nitrogen levels as well as with nodule dry mass. Two *B. diazoefficiens* strains stood out and possessed *nosZ*. In contrast, *B. elkanii* and *B. japonicum* isolates displayed lower chlorophyll and symbiotic performance, and lacked *nosZ*. Our findings highlight the potential of measuring chlorophyll contents and testing for the presence of *nosZ* as two straightforward techniques that may enhance quality control, enabling selection of superior and safe locally isolated strains for soybean inoculants.

## 1. Introduction

Global warming is becoming evident year after year, whereby it is urgent to strengthen measures to mitigate greenhouse gas emissions. Among the most contaminant human activities is the irrational use of N-fertilizers, which, being applied in up to 10-fold excess in several crops, leach into the soil and decompose producing N_2_O gas with strong greenhouse and global warming effects (Erisman *et al*., 2013; Aryal *et al*., 2022). Therefore, measures intended to mitigate the extended use of N-fertilizers are urgent. It is well known that biological N_2_ fixation may replace N-fertilizers in legume crops, and even incorporate N to the soil (Imran *et al*., 2021). Soybean [*Glycine max* (L.) Merr.], the most important grain legume worldwide, is a host for diverse N_2_-fixing bacteria and, in particular, *Bradyrhizobium diazoefficiens*, *B. elkanii*, and *B. japonicum* are commonly used as inoculants in vast cropping areas. However, mineralized N from decomposing roots, nodules, and aerial biomass incorporated as green manure to the soil may undergo denitrification by soil microbiota and release N_2_O if the pathway is not completed until total reduction to N_2_. It was reported that inoculation with certain strains of *Bradyrhizobium* spp. may produce N_2_O due to incomplete denitrification, while others are capable of completing this pathway, depending on whether they possess the N_2_O reductase encoded in the *nosZ* gene (Siqueira *et al*., 2017; Obando *et al*., 2019; Mania *et al*., 2020). Therefore, inoculation of soybean with *nosZ*^+^ strains leaded to a significant reduction of N_2_O emission from this crop in comparison with plots inoculated with *nosZ*^‒^ strains (Akiyama *et al*., 2016).

In addition, for N_2_ fixation to be efficient, a positive net balance between N incorporated by the plant from atmosphere and from soil must be achieved. In Argentina, the third-largest producer of soybeans, it was estimated that soybean fixes 2.54 Tg of N per growth cycle, contributing 17% of total symbiotic N_2_ fixation globally. Since, in the same conditions, 2.43 Tg N are exported within seeds at harvest, it was estimated that the net average gain of N by this crop system is 0.11 Tg (Collino *et al*., 2015). However, this balance depends on various factors, including environmental conditions, plant cultivars, and the prevalence of *Bradyrhizobium* spp. genotypes in the soil. Although soybean crops in Argentina are widely inoculated with the elite strain *B. japonicum* E109 (Torres *et al*., 2015), other strains are prevalent inside nodules, suggesting a poor adaptability of E109 to the conditions of agricultural soils in Argentina (Iturralde *et al*., 2019). In addition, E109 is known as a *nosZ*^‒^ strain (Obando *et al*., 2019) whereby its use as a universal inoculant in Argentina may be harmful regarding N_2_O emissions. Therefore, it would be of interest to measure and register the N_2_-fixing potential and the presence of *nosZ* in *Bradyrhizobium* spp. strains prevalent in a defined crop area.

Several methods for estimating N_2_-fixation are known from long ago (Burris, 1972), but many are cumbersome, difficult to apply in non-destructive or field conditions, or require costly equipment. One popular method is the acetylene reduction assay, but it was criticized because exposure of nodulated roots to acetylene causes a rapid decline in nitrogenase activity, which was attributed to the cessation of ammonia production by nitrogenase and increase of oxygen diffusion resistance by the nodules (Minchin and Witty, 1989). Furthermore, the magnitude of this decline depends on environmental conditions, rhizobial species or strains, whereby this method is not recommended for comparative studies. Another method involves measuring H_2_ evolution, which is a by-product of the nitrogenase reaction (Witty and Minchin, 1998), but it is restricted to strains lacking the ability to recycle H_2_ for generation of extra ATP (Carter *et al*., 1978). Other methods involve direct measurement of total N accumulated by plants growing hydroponically without added combined N or the measurement of N_2_-fixation export products, such as ureides in the case of soybean (Herridge, 1982). However, these methods are laborious and often destructive, limiting their application in serial measurements or selection procedures involving individual plants.

Alternative methods, such as the use of the ^15^N isotope, can be applied in field trials but are also destructive and costly (Chalk and Ladha, 1999). Thus, perhaps due to unavailability of rapid, nondestructive, simple, and reliable tests for N_2_-fixation, the inoculants industry seldom measures this important symbiotic parameter as part of the quality control of its strains (Penna *et al*., 2011; Herrmann and Lesueur, 2013; Herridge *et al*., 2014; de Souza *et al*., 2019).

As an alternative, indirect estimation of N_2_-fixation could be achieved by measuring parameters related to photosynthesis, as it is known that the N-status of leaves is correlated with the rate of photosynthesis in different plant species. The majority of N allocated to leaves is integrated into the photosynthetic apparatus (Evans and Clarke, 2019). Therefore, gross leaf photosynthesis and CO_2_ uptake rate are positively correlated with leaf N concentration in soybean (Boote *et al*., 1978; Lugg and Sinclair, 1981; Boon-Long *et al*., 1983; Buttery and Buzzell, 1988). However, other experiments have shown no response in photosynthesis rate to different levels of N-fertilization in soybean (Moreira *et al*., 2015). The presence of the N_2_-fixing bacteria may have masked the responses to varying N-fertilization levels. Furthermore, a positive correlation between chlorophyll and leaf N content has been observed in field studies, although it was not strong enough to be used as a predictor of leaf N content (Fritschi and Ray, 2007). Nevertheless, the nodulation state of these plants was not reported by the authors.

The above studies have mostly focused on soybean plants in the pod-filling state, and photosynthesis rate and leaf N content may differ in earlier vegetative stages due to factors such as changes in leaf mass per unit area. In soybean plants grown hydroponically for 23-30 days after germination (around V2 stage), a significant positive correlation was observed between N content per unit leaf area and light-saturated photosynthesis (Adams *et al*., 2018). However, the nodulation state of these plants again was not reported. Therefore, it remains unclear whether photosynthesis-related parameters can be used as predictors of N_2_-fixation by root nodule bacteria in soybean.

A non-destructive and rapid method for estimating photosynthesis rate in soybean is the measurement of leaf chlorophyll content using a portable chlorophyllometer (Ma *et al*., 1995; Castelli *et al*., 1996). This instrument measures leaf transmittance in the red and infrared wavelengths, allowing for determination of relative chlorophyll content spectrophotometrically using the infrared measure as a self-correction. Although the relationship between chlorophyll content and chlorophyllometer readings is non-linear (Markwell *et al*., 1995; Uddling *et al*., 2007), the measurements are being interpreted as a reflection of a plant’s nutritional status in agronomic research. However, previous investigations have failed to consider the role of nodulation in this relationship.

Since the use of a chlorophyllometer might have potential to become a simple and nondestructive routine test of N_2_-fixation, in this report we conducted measurements of various symbiotic parameters and chlorophyll relative contents as determined with a chlorophyllometer in soybean plants nodulated by different strains of *Bradyrhizobium* spp. isolated from an agricultural field at the Province of Buenos Aires, Argentina, in order to assess the correlation among these measures. We found that the chlorophyllometer readings had a good predictive power for the N_2_- fixing quality of the different strains. In addition, we designed primers to detect the presence of *nosZ* by PCR in these strains. These approaches could be incorporated into quality control protocols for strains assessment in the production of safe and efficient inoculants and supports the recommendation of new strains adapted to local conditions.

## 2. Materials and Methods

### 2.1. Bacterial strains, plants, and culture conditions

We obtained *B. diazoefficiens* USDA 110^T^ from Deane Weber, ARS-USDA, United States, *B. japonicum* E109 from Alejandro Perticari, IMYZA-INTA, Argentina, *B. elkanii* USDA 76^T^, and *B. japonicum* USDA 6^T^ from Esperanza Martínez-Romero, Centro de Ciencias Genómicas, UNAM, México, *B. diazoefficiens* SEMIA 5080, and *B. japonicum* SEMIA 5079 from Mariangela Hungria, EMBRAPA, Brazil, and *Azospirillum argentinense* Az39^T^ from Fabricio Cassán, UNRC, Argentina. We maintained all these strains, as well as the *Bradyrhizobium* spp. isolated in this study (see section 3.1) in glycerol stocks as described (Iturralde *et al*., 2019). For routine stocks, we grew the bacteria in YMA (Vincent, 1970), and for plants inoculation we grew them in a liquid medium that contained (in g l^‒1^ distilled water): glucose, 5; mannitol, 5; sodium gluconate, 5; yeast extract, 2.2; CaCl_2_, 0.9; MgCl_2_, 0.3; K_2_HPO_4_, 1.1 and KH_2_PO_4_, 0.9. Soybean Bioceres 4.12 RR was provided by Bioceres S.A., Argentina. We surface-sterilized and germinated the seeds as already described (Iturralde *et al*., 2019). After germination, we transferred plantlets with 3 cm-long roots to 500-ml plastic pots filled with vermiculite and watered with 200 ml of N-free modified Fåhraeus nutrient solution (MFS, Lodeiro *et al*., 2000). Then, we cultivated these plants in a greenhouse at 30 °C/20 °C day/night temperature with light supplemented in order to obtain 16 h photophase for the periods indicated for each experiment. We watered the plants two times per week (once with sterile MFS and once with sterile distilled water).

### 2.2. Isolation of soybean-nodulating rhizobia

We obtained soil samples in January 2021 from an experimental field of Facultad de Ciencias Agrarias y Forestales (Faculty of Agricultural and Forestry Sciences), UNLP, located at Los Hornos, Province of Buenos Aires, Argentina (34°59’10.5” S; 58°59’58.9” W), and from the garden of one of the authors (ARL) located at Mar del Plata, Province of Buenos Aires, Argentina (38°4’ 23.0” S; 57°33’34.5” W). The Los Hornos soil was under experimentation conducted by the Faculty of Agricultural and Forestry Sciences to evaluate different soybean inoculants and other crops systems since at least the previous 10 years before sampling. The Mar del Plata soil has no previous history of soybean. From 5 sites of each location, we collected soil samples at the vertices of a zigzag and in the middle of the plot and stored the samples in plastic bags. All samples were kept refrigerated and transported to the laboratory as soon as possible. To obtain the soybean-nodulating population from both soil samples, we extracted the bacteria from soils by suspending 100 g soil in 1 l MFS and agitated the suspension for 1 h at 180 rpm in an orbital shaker. Then, we let the soil to settle, and centrifuged the supernatant at 3,000 × *g* 5 min to obtain the bacterial samples. With these suspensions we inoculated 5 soybean plants and cultivated them for 21 days until nodules developed. As controls, we inoculated 5 plants with *B. diazoefficiens* USDA 110 and another 5 plants with MFS without bacteria. After checking that the positive controls developed normal nodules and the negative controls presented no nodules, we collected the nodules produced by plants inoculated with each soil extract. We surface-sterilized and crushed all these nodules and afterwards streaked their contents in YMA as described (Iturralde *et al*. 2019). We purified the isolates by colony isolation and picking three consecutive times and then we passed the bacteria through nodules and reisolated them again. Finally, we analyzed their genotypes by DNA fingerprint and determined the species by gene sequencing (see section 2.3). We denominated the isolates with two letters that indicate the site of origin, a letter “S” that indicates that the isolate comes from a soil sample, and a number that identifies the isolate. For instance, the denomination “LH/S-10” names the isolate #10, which was obtained from a soil sample in Los Hornos, and “MP/S-01” names the isolate #1 which was obtained from a soil sample in Mar del Plata.

### 2.3. DNA extraction, gene sequencing and genotypic analysis

We isolated genomic DNA from each individual bacterial strain as described (Iturralde *et al*. 2019). We used the same PCR conditions and primers already described for DNA-fingerprint and DNA amplification of 16S rRNA gene (V1-V9), *atpD*, *glnII* and *recA* fragments (Iturralde *et al*., 2019). For DNA-fingerprint analysis, we separated the BOX-A1R PCR products in agarose (2% w/v) gel electrophoresis at 70 volts during 3 h. We normalized the gel photographs with reference lanes consisting of molecular weight standards (DNA ladder plus 100 bp, Genbiotech, Buenos Aires, Argentina) and reaction products with DNA from *B. diazoefficiens* USDA 110. Then, we analyzed each bacteria population lane for the presence or absence of bands with the GelCompare II 4.0 software (Applied Maths, Kortrijk, Belgium). We optimized the band pattern with a 1.5% optimization and 1.7% tolerance. We obtained the cladogram with the unweighted pair-group method with arithmetic mean (UPGMA) (Sneath and Sokal, 1973) and evaluated the distances among branches with the Jaccard coefficient (Jaccard, 1912). For DNA sequencing, we obtained DNA fragments corresponding to 16S rRNA, *atpD*, *glnII*, and *recA* genes by PCR using the primers and amplification conditions already described (Iturralde *et al*., 2019). Then, we send the DNA fragments to Macrogen (Korea) for their sequencing with an ABI 3730xl DNA sequencer. We deposited the obtained sequences in GenBank with the accession numbers detailed in Supplementary Table S1. We analyzed the 16S rRNA and *atpD-glnII-recA* genes with DNASTAR Lasergene (DNAStar, Inc.) and compared these sequences with related sequences from GenBank with NCBI BLAST. Then, we aligned the sequences with MUSCLE and constructed phylogenetic trees using MEGA 11 with neighbor joining method (Tamura *et al*., 2021). For analysis of DNA sequence data, we selected the DNA evolution model using the above-mentioned software.

Furthermore, we used the Tamura 3-parameter model with gamma distribution (T92+G) to obtain the cladograms for 16S rRNA and concatenated *atpD-glnII-recA* genes. We grouped the sequences by maximum likelihood and evaluated the support for tree nodes by bootstrap analyses with 1,000 samplings (Felsenstein, 1985; Hedges, 1992). For *nosZ* detection, an internal 200-bp fragment from this gene was obtained by PCR using the primers and cycle conditions already described (Itakura *et al*., 2013).

### 2.4. Analysis of plant growth-promoting activities

We assayed phosphate solubilization by measuring the ratio of the diameter of the solubilization halo to the diameter of the colony in NBRIP agar medium (Nautiyal, 1999). For siderophore production we cultured the strains in CAS (chromium azurol S) agar medium (Schwyn and Neilands 1987) and detected siderophore production by the overlay method described by Pérez-Miranda *et al*. (2007). For detection and quantification of indoleacetic acid (IAA) production we centrifuged YM-grown bacteria at 8,220 × *g* for 20 min and filtered the supernatant through 0.22 μm cellulose filters (Millipore, USA). Then, we quantified IAA in the filtered supernatants by high-performance liquid chromatography (HPLC) using purified IAA (Sigma-Aldrich, Buenos Aires, Argentina) as standard, according to Jensen *et al*., (1995).

### 2.5. Microscopy

Bacterial cells were heat-fixed to slides and Gram staining was carried out with crystal violet and safranin. The slides were washed and dried for optical microscopy. Isolated cells were observed at 1000 × using a Nikon Eclipse E400 microscope.

### 2.6. Plant assays

To determine the most probable numbers of rhizobia in each soil sample we proceeded according to Vincent (1970). We prepared serial 1/2 dilutions from each soil extract and inoculated 5 plants with each of these dilutions. Then, we recorded the number of nodulated plants with each dilution and deduced the most probable number using appropriate tables (Vincent, 1970). To inoculate the plants for symbiotic studies, we suspended single bacterial colonies from YMA plates in 1 ml sterile MFS and deposited this suspension onto each seedling after planting in the vermiculite pots. Seven seedlings per strain were inoculated. As controls, we inoculated a set of plants with 1 ml of sterile MFS without bacteria, and as reference we used *B. japonicum* E109 inoculated in separate pots. Then, we cultured the plants for 28 days in the conditions already described. After this growth period, we separated the shoot, the root, and the nodules from each plant. We counted the total number of nodules of each plant, and then we dried all the materials at 60°C up to weight constancy. For determination of total N, we sent the dried materials to C & D Laboratory (Buenos Aires, Argentina) for Kjeldahl analysis.

### 2.7. Estimation of chlorophyll contents

Leaf chlorophyll contents were estimated with a SPAD-502 chlorophyllometer (Minolta, Japan) reading each leaflet of a trifoliate leaf at V1 and V2 stages. The readings of each leaflet were combined, and the results were expressed in relative soil-plant analysis development (SPAD) units as the average for each trifoliate leaf.

### 2.8. Statistical analysis

For principal component analysis (PCA) we used InfoStat software (Di Rienzo et al., 2009) and for the symbiotic parameters we employed the analysis of variance (ANOVA) followed by Tukey test to identify significant mean differences with a threshold of *p*< 0.05 as already described (Iturralde *et al*., 2019). We estimated the correlations between the different symbiotic variables and chlorophyll contents by Pearson correlation using Excel. The significance of the correlation coefficients obtained was calculated using the Student’s *t*-test.

## 3. Results

### 3.1. Genotypic analysis and classification

We obtained 21isolates from soil extracts using soybean as trap-plant: 16 isolates from Los Hornos and 5 isolates from Mar del Plata, although this last soil contained much less rhizobia (most probable numbers were 7.6·10^3^ rhizobia·g^-1^soil in Los Hornos, and 6.0·10^1^ rhizobia·g^-1^ soil in Mar del Plata). All isolates grew in YMA supplemented with Congo Red, producing slow-growth, white mucous colonies with smooth borders, typical of *Bradyrhizobium* spp. In addition, all isolates were Gram-negative rods. Genetic diversity was analyzed through BOX-A1R DNA-fingerprints (Supplementary Fig. S1) and a dendrogram, based on UPGMA algorithm and Dice coefficient was constructed using *B. diazoefficiens* USDA 110^T^, *B. japonicum* E109, *B. elkanii* USDA 76^T^, *B. japonicum* USDA 6^T^, *B. diazoefficiens* SEMIA 5080, and *B. japonicum* SEMIA 5079 as reference strains (Fig. 1). By using a 40% similarity cutoff we grouped the isolates in three main clades.

**Fig. 1.**
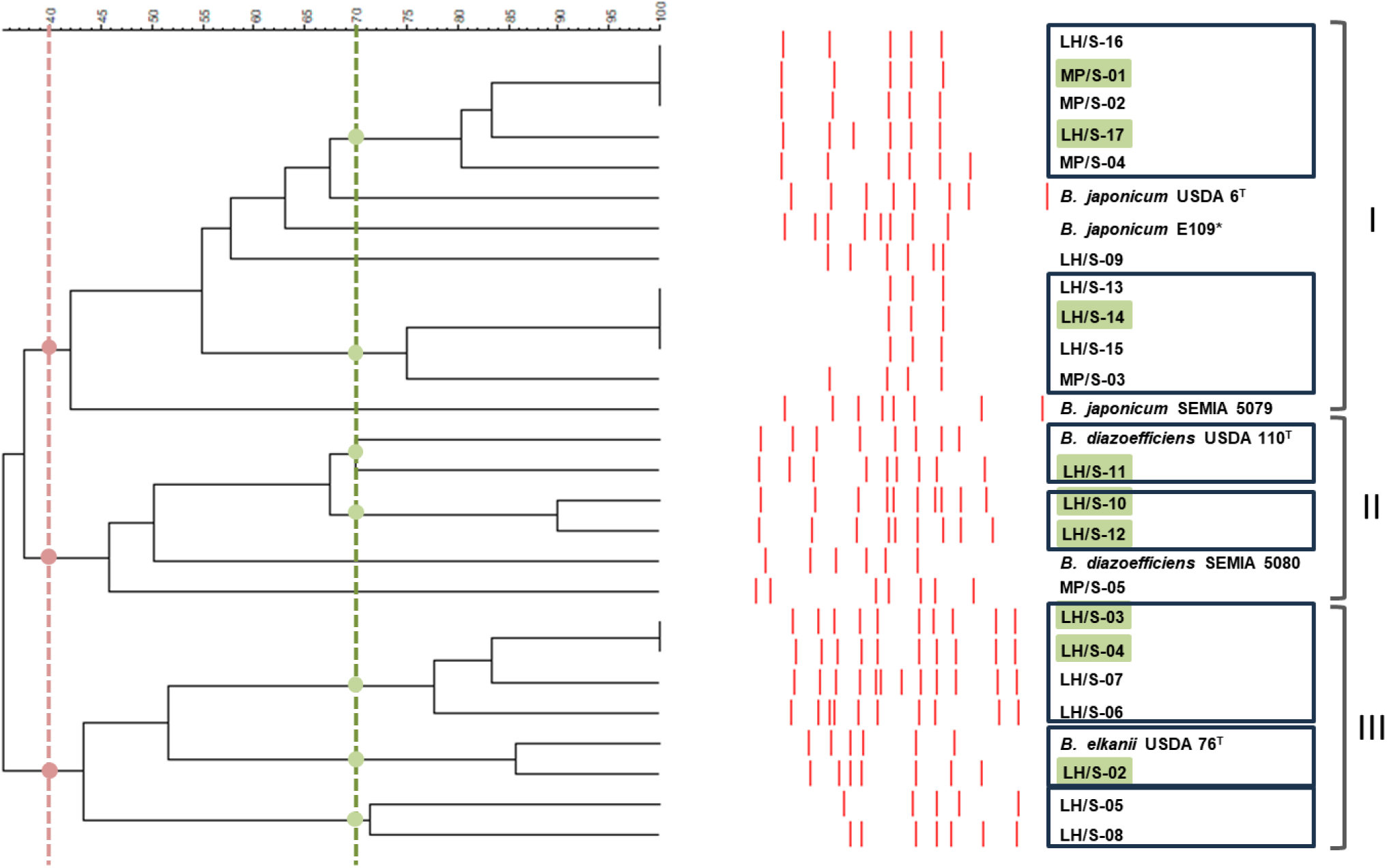
DNA-fingerprint of the soybean-nodulating isolates and reference strains obtained with BOX-A1R primer. Clades (numbered at the right) were defined at 40% similarity (red lines, dots, and brackets). The isolates with more than 70% similarity are presumed as sharing the same parental genotype (boxed). Isolates denoted with a shadowed box are those picked for further species identification and characterization. T indicates type strains; the asterisk indicates the reference strain.

Clade I contained 10 isolates and the three *B. japonicum* reference strains; clade II contained 4 isolates along with the two *B. diazoefficiens* reference strains, and clade III grouped the remaining 7 isolates together with the *B. elkanii* reference strain. In addition, we analyzed the distribution of the isolates using a 70% cutoff, which was considered that may group genotypes sharing the same parental genotype in *Bradyrhizobium* spp. (Loureiro *et al*., 2007). This analysis rendered 7 groups of isolates plus 2 isolates that remained alone. Only one isolate was similar to *B. diazoefficiens* USDA 110 and another to *B. elkanii* USDA 76 according to this criterion, while no isolates displayed substantial similarity to *B. japonicum* E109 (Fig. 1).

Subsequently, we selected 9 isolates as representative of the three main clades for species determination and *nosZ* detection. Eight isolates were from Los Hornos showing differences in the DNA fingerprint banding pattern and one isolate was from Mar del Plata.

We sequenced the 16S rRNA genes from these 9 isolates in order to achieve a taxonomic approximation. We observed that isolates LH/S-02, LH/S-03 and LH/S-04 showed a close relationship with several *B. elkanii* strains as well as *B. jicamae* PAC68^T^, *B. valentinum* LmjM3, *B. icense* LMTR 13^T^, *B. embrapense* SEMIA 6208^T^ and *B. lablabi* CCBAU 23086^T^, while LH/S-10, LH/S-11 and LH/S-12 were close to *B. diazoefficiens* SEMIA5080 and USDA110^T^ strains, and LH/S-14, LH/S-17 and MP/S-01 were close to *B. japonicum* USDA6^T^, SEMIA5079 and E109 strains (Fig. 2). These relationships between isolates and reference species were in close agreement with those observed in the DNA fingerprint.

**Fig. 2.**
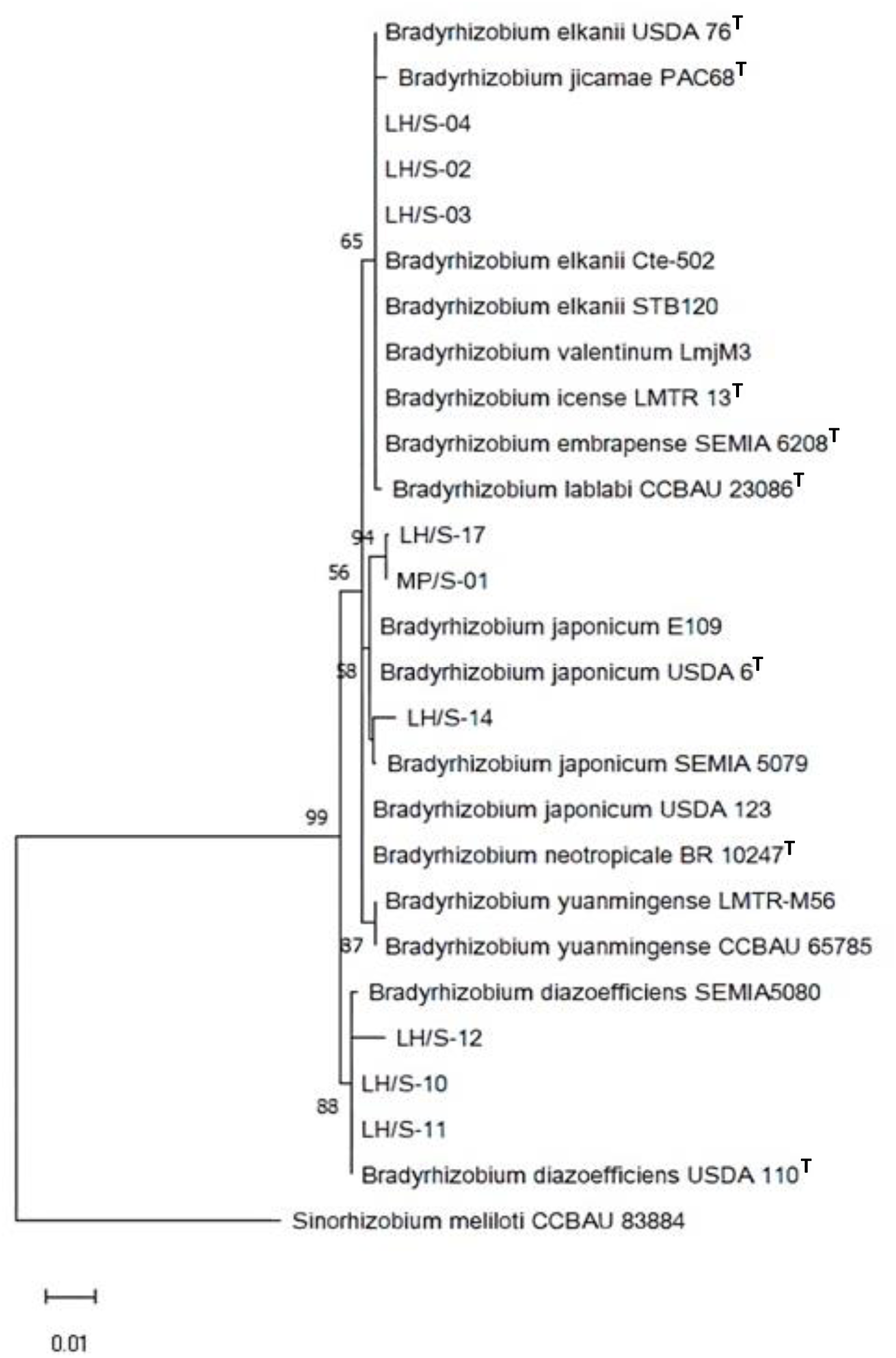
Maximum likelihood cladogram obtained from 16S rRNA gene of soybean-nodulating isolates and type/reference strains. Bootstrap values greater than 50% calculated for 1000 replications are indicated at internodes. Branch lengths are shown to scale, indicating evolutionary distance used to infer the phylogenetic tree. *Sinorhizobium meliloti* CCBAU 83884 was included as outgroup. T indicates type strain.

Since 16S ribosomal RNA gene sequences lack sufficient information to discriminate between *B. diazoefficiens* and *B. japonicum*, we compared the concatenated *recA-atpD-glnII* sequences to refine the phylogenetic analysis (Menna *et al*., 2009; Delamuta *et al*., 2013). We performed the multilocus sequence analysis by considering only the complete aligned sequences of the soybean-nodulating strains and the type/reference strains sequences (size in parentheses) retrieved from GenBank: 16S rRNA (1,135 bp), *recA* (434 bp), *atpD* (356 bp), *glnII* (282 bp). This analysis confirmed the results obtained with 16S rRNA sequence comparisons and BOX-AR1 DNA fingerprinting, indicating that three of the *Bradyrhizobium* spp. isolates were related to *B. japonicum*, three to *B. diazoefficiens*, and the remaining three to *B. elkanii* (Fig. 3).

**Fig. 3.**
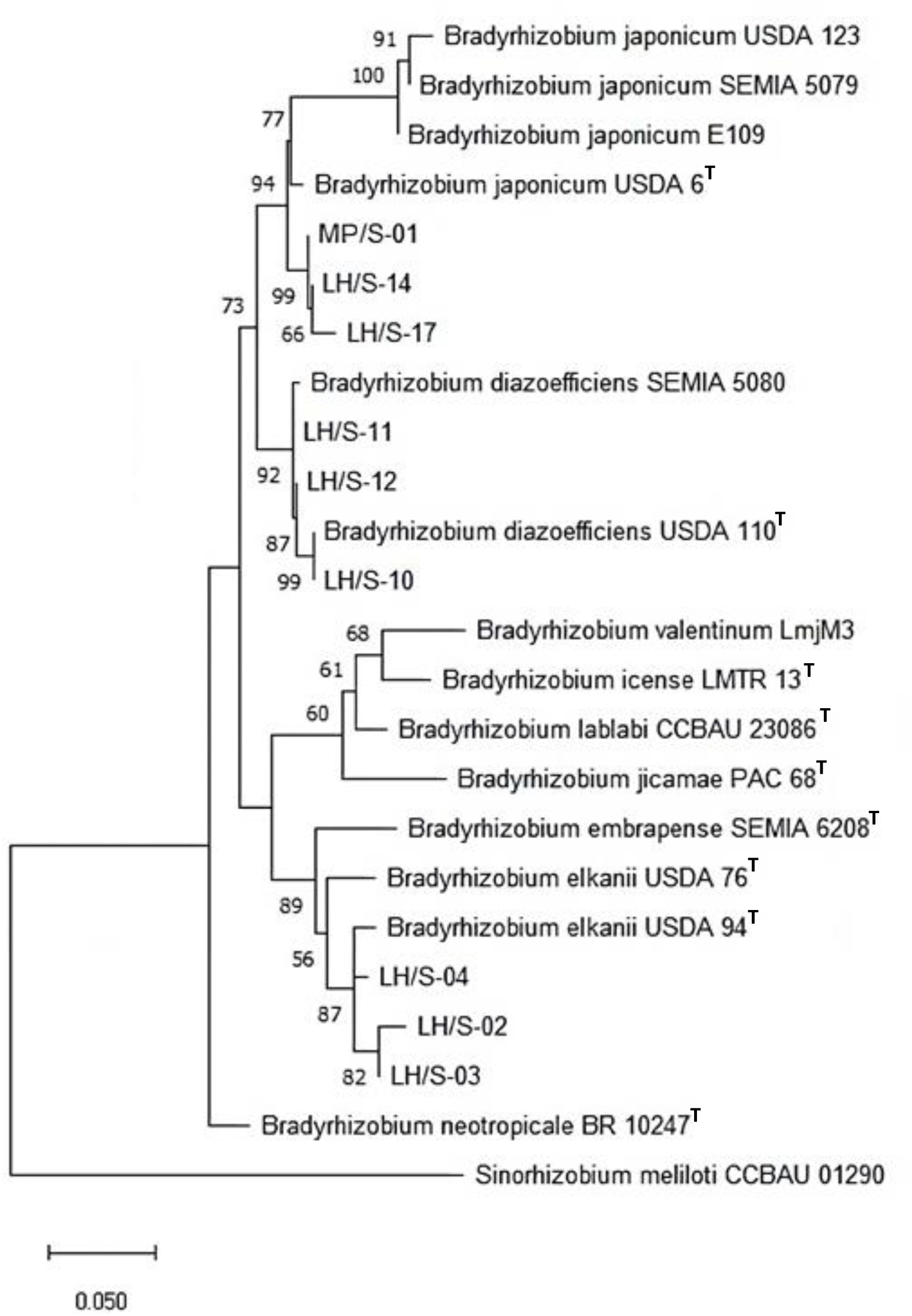
Maximum likelihood cladogram obtained for the concatenated *atpD*-*glnII*-*recA* fragment sequences to distinguish *B. diazoefficiens*, *B. japonicum* and *B. elkanii* among picked soybean- nodulating isolates in comparison with type/reference strains. Bootstrap values greater than 50% calculated for 1000 replications are indicated at internodes. Branch lengths are shown to scale, indicating evolutionary distance used to infer the phylogenetic tree. *Sinorhizobium meliloti* CCBAU 01290 was included as outgroup. T indicates type strain.

To detect the *nosZ* gene that encodes the N_2_O reductase we used two sets of primers: one specific for *nosZ* and the other specific for the housekeeping gene *recA* as positive control for the integrity of DNA templates. Our PCR analysis indicated that, as reported previously, *B. diazoefficiens* USDA 110 is *nosZ*^+^ since the target 200-bp *nosZ* fragment was amplified, while *B. japonicum* E109 is *nosZ*^‒^. These results are in agreement with the complete genomic sequences of these strains (Kaneko *et al*., 2002; Torres *et al*., 2015). In turn, the isolates were also diverse in this respect: while the three *B. diazoefficiens* isolates were *nosZ*^+^, all *B. japonicum* and *B. elkanii* were *nosZ*^‒^ (Fig. 4).

**Fig. 4.**
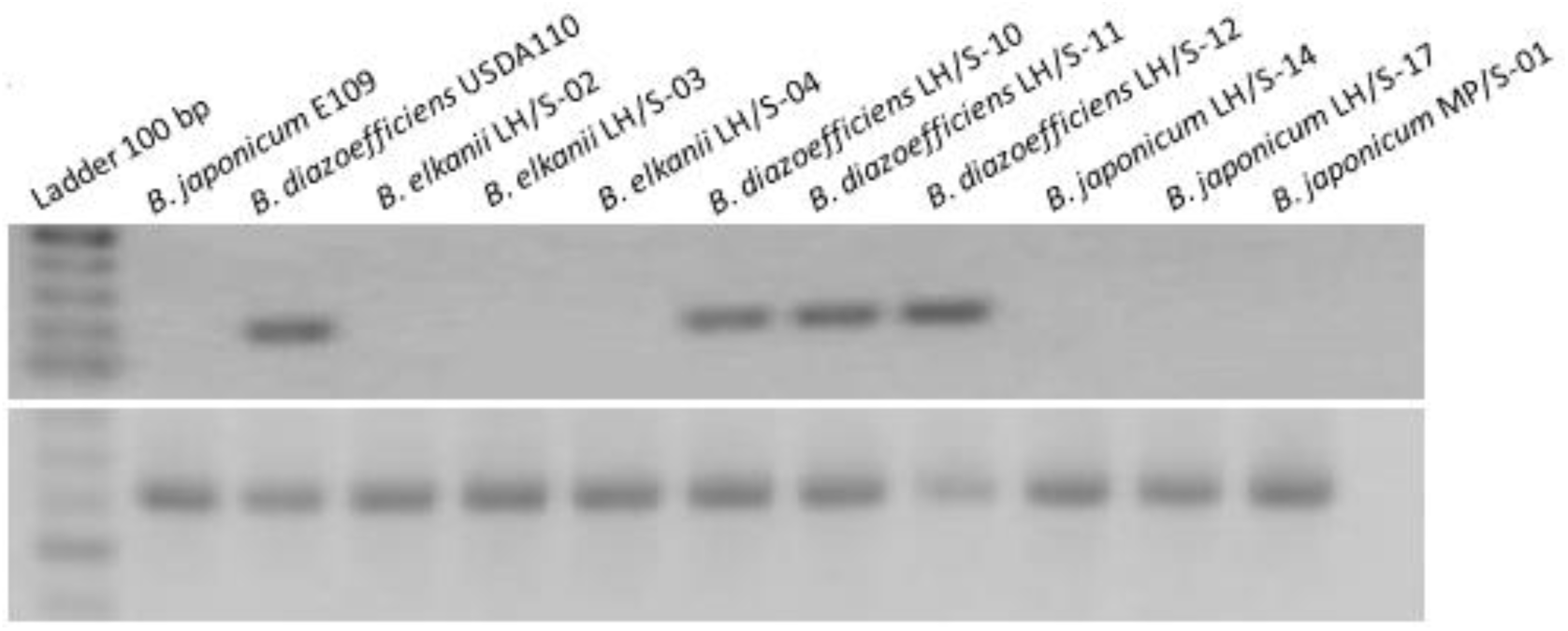
Detection of the 200-bp amplicon of *nosZ* by PCR in the soybean-nodulating field isolates (upper panel). As internal control for the integrity of the DNA templates, the housekeeping *recA* 650-bp amplicon was obtained (lower panel).

### 3.2. Plant growth-promoting activities

None of the 9 selected isolates were able to solubilize phosphate or produce siderophores. To evaluate the IAA production and secretion *in vitro*, we measured the concentration of IAA in culture supernatants by HPLC using purified IAA as standard, and the IAA-producing strain *Azospirillum argentinense* Az-39 (Molina *et al*., 2018, Dos Santos *et al*., 2022) as positive control (Supplementary Fig. S2). The isolate *B. elkanii* LH/S-02 secreted 1.9 ± 0.86 μg · ml^‒1^, *B. elkanii* LH/S-03 secreted 1.66 ± 0.39 μg · ml^‒1^ and *B. elkanii* LH/S-04 secreted 0.55 ± 0.02 μg · ml^‒1^ IAA. By comparison, *A. argentinense* Az39 secreted 4.95± 1.29 μg · ml^‒1^ IAA in the same conditions.

None of the other isolates were able to secrete IAA in detectable amounts. These results indicated that, among the isolates analyzed, production and secretion of IAA was a property of *B. elkanii*, not shared with *B. diazoefficiens* or *B. japonicum*.

### 3.3. Symbiotic interaction between the selected isolates and soybean

To assess the symbiotic proficiency of our selected isolates we conducted a series of experiments in the greenhouse, where individual plants were inoculated with each isolate, a set of plants was inoculated with *B. japonicum* E109 as reference strain, and another set of plants was kept uninoculated. We measured the following parameters: numbers of nodules per plant, individual nodule dry weight, shoot and root fresh and dry weights, shoot/root fresh and dry weight ratios, N contents per biomass unit in dry shoots, and chlorophyll contents in leaves. The detailed results are presented in Supplementary Table S2 and here we will focus on the most relevant relationships.

To include all variables mentioned and their relationships, we performed a principal component analysis (PCA). As shown in Fig. 5, the principal component (PC) 1 explained 49.1% of total variance, while the PC 2 explained 32.7%. Thus, both components rescued 81.8% of variance. The stronger correlations may be observed among the symbiotic variables individual nodule dry weight, number of nodules per plant, and N contents per biomass unit, and among the biomass variables shoot dry weight and root dry weight. Both groups of variables point in the direction of PC 1, where chlorophyll contents and shoot fresh weight have the highest relationship. On the other hand, shoot/root ratio (dry) is not correlated with shoot dry weight and root dry weight, but it has correlation with the symbiotic variables individual nodule dry weight, number of nodules per plant and N contents per biomass unit. There was also no correlation between shoot fresh weight and root fresh weight, and, in addition, shoot/root ratio (fresh) was important for both PC 1 and PC 2, but nearly opposite to shoot fresh weight and root fresh weight (Fig. 5).

**Fig. 5.**
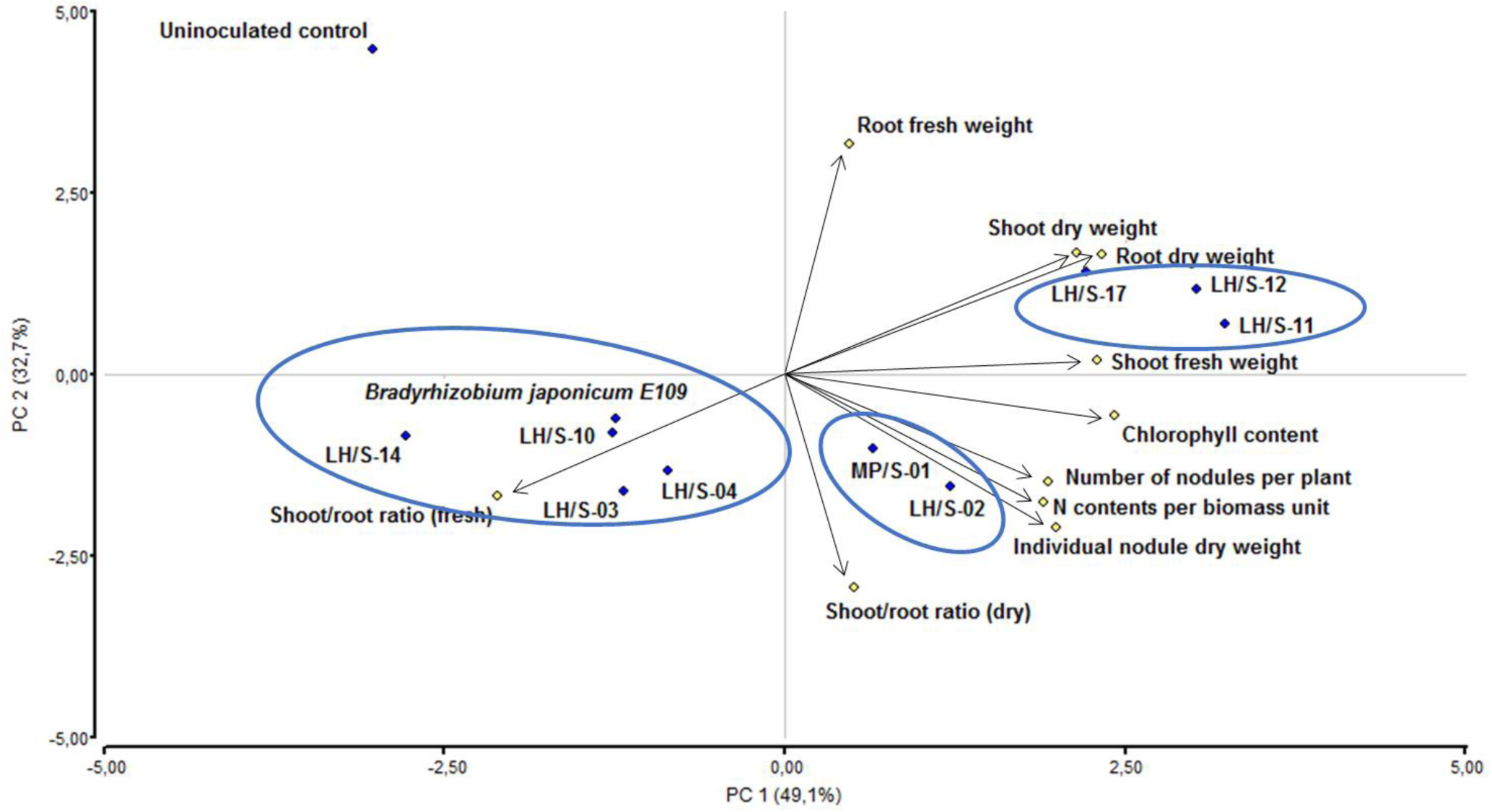
Principal components analysis (PCA) of number of nodules per plant, individual nodule dry weight, shoot and root fresh weight, shoot and root dry weight, shoot/root fresh and dry weight ratios, N contents per biomass unit in dry shoots, and chlorophyll contents in leaves of soybean plants inoculated with the field soybean-nodulating isolates and *B. japonicum* E109 as reference strain. Principal components PC 1 and PC 2 explained 49.1% and 32.7%, respectively, of the total variance.

The isolates formed three groups: one composed by *B. elkanii* LH/S-03 and LH/S-04, *B. japonicum* LH/S-14 and E109, and *B. diazoefficiens* LH/S-10, which was characterized by shoot/root ratio (fresh), another composed by *B. diazoefficiens* LH/S-11 and LH/S-12, and *B. japonicum* LH/S-17, which was characterized by the biomass variables shoot dry weight, root dry weight and shoot fresh weight, and the third one composed by *B. elkanii* LH/S-02 and *B. japonicum* MP/S-01, which was characterized by the symbiotic variables individual nodule dry weight, number of nodules per plant and N contents per biomass unit. These last two groups were also characterized by chlorophyll contents (Fig. 5).

We calculated the total N-contents of shoots (hereafter referred to as shoot N) from N contents per biomass unit and shoot dry weight and expressed the results in mg·N plant^‒1^, the total nodules dry weight by multiplying the number of nodules per plant by individual nodule dry weight and expressed it in mg·plant^‒1^, and finally we expressed chlorophyll contents in SPAD units. Since these plants did not receive N in the plant nutrient solution, shoot N reflects the N obtained from atmosphere, while total nodules dry weight is an indicator of nodular activity. As shown in Fig. 6 A-C, we observed a wide diversity of chlorophyll contents, shoot N, and total nodules dry weight among the plants inoculated with the different isolates, indicating substantial differences in photosynthetic and N_2_-fixation capacities. This result allowed us to study the possible correlation between each of the symbiotic variables with photosynthesis, evaluated through chlorophyll contents. As shown in Fig. 6 D,E we indeed observed statistical significant correlations between the pairs of variables analyzed. For the pair shoot N vs chlorophyll contents the Pearson correlation coefficient was 0.81 and for total nodules dry weight vs chlorophyll contents it was 0.61 (both significant with *p* < 0.005).

**Fig. 6.**
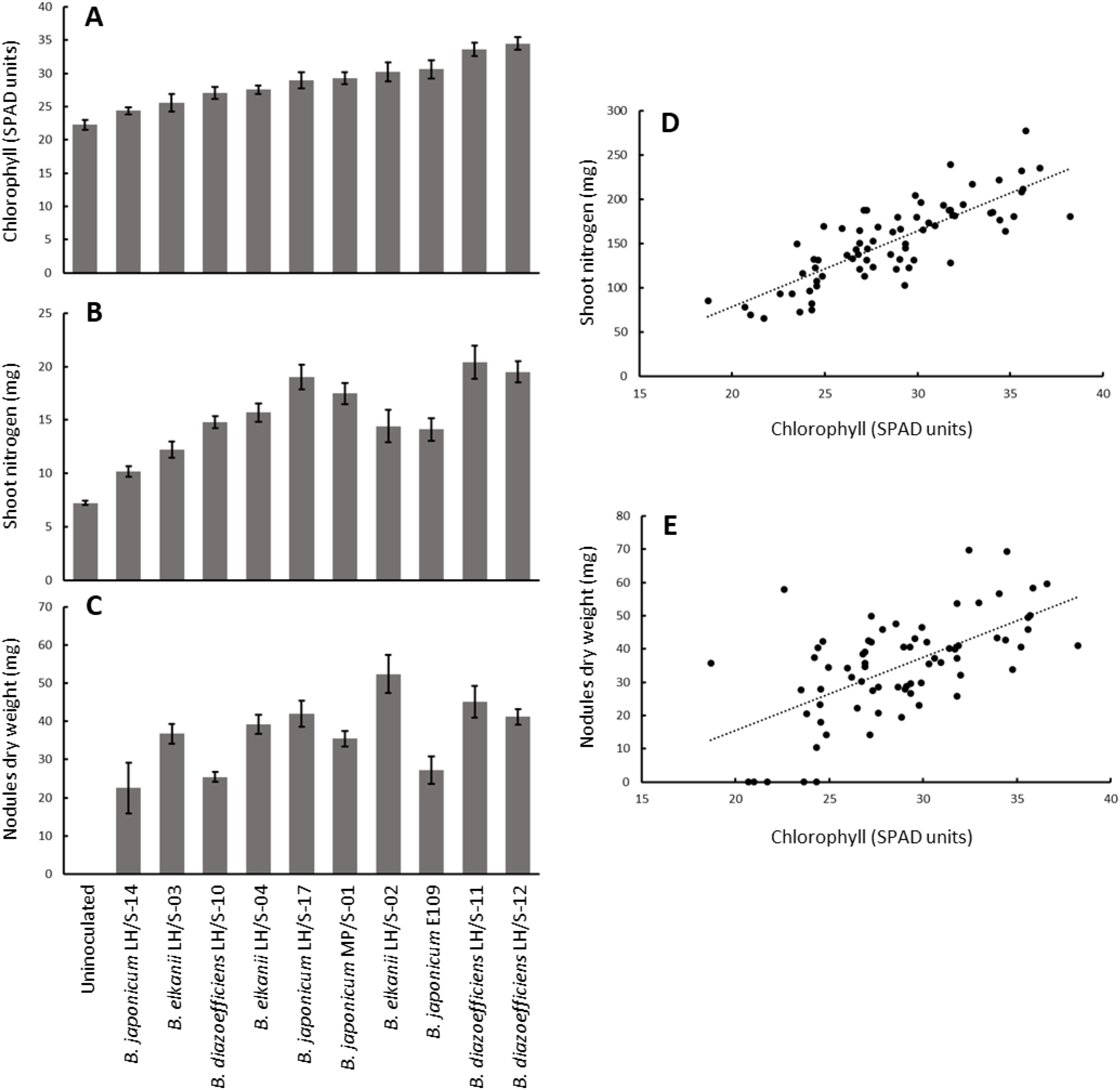
Diversity of symbiotic-related values and their correlations with chlorophyll contents in soybeans inoculated with the different soybean-nodulating field isolates, *B. japonicum* E109 as reference strain, and an uninoculated control. (**A**) Chlorophyll contents in leaves. (**B**) Shoot nitrogen accumulated per plant. (**C**) Total nodules dry weight per plant. (**D**) Correlation between shoot nitrogen and chlorophyll contents. (**E**) Correlation between total nodules dry weight and chlorophyll contents. Error bars in **A**-**C** are standard errors of the mean. In **D**, *r* = 0.81. In **E**, *r* = 0.61. Both correlation coefficients were significant with *p*< 0.005.

If we order the isolates in thirds from higher to lower value (excluding *B. japonicum* E109), we can appreciate that the top third as regards shoot N is integrated by *B. diazoefficiens* LH/S-11 (1^st^), *B. diazoefficiens* LH/S-12 (2^nd^) and *B. japonicum* LH/S-17 (3^rd^), while the top third as regards total nodules dry weight contains *B. elkanii* LH/S-02 (1^st^), *B. diazoefficiens* LH/S-11 (2^nd^), and *B. diazoefficiens* LH/S-12 (3^rd^). On the other hand, the top third as regards chlorophyll contents is composed by *B. diazoefficiens* LH/S-12 (1^st^), *B. diazoefficiens* LH/S-11 (2^nd^), and *B. elkanii* LH/S- 02 (3^rd^). By comparing with the PCA in Fig. 5, we can see that the isolates LH/S-11, LH/S-12, and LH/S-17 are situated in the same region, characterized by biomass production variables, while LH/S-02 is characterized by symbiotic variables. Hence, the measurement of chlorophyll contents using the chlorophyllometer correctly identified *B. diazoefficiens* LH/S-11 and *B. diazoefficiens* LH/S-12 as the most prominent isolates regarding their symbiotic potential among the 9 isolates analyzed.

## 4. Discussion

By using soybean as trap-plant we obtained 21 isolates from soil samples taken at Los Hornos (LH) and Mar del Plata (MP), two locations separated by 370 km in the Province of Buenos Aires, Argentina. Soybean was cultivated in LH for experimental purposes since the last 10 years, while the MP samples come from a garden where soybean was never planted. By using BOX-AR1 DNA fingerprint we grouped these isolates in three genotypic clades, each one represented by reference strains of *B. japonicum* (clade I), *B. diazoefficiens* (clade II) and *B. elkanii* (clade III). Then, we determined the species to which 9 selected isolates belong by sequencing the 16S rRNA, *recA*, *atpD*, and *glnII* genes. By comparing this information with the genotypical distribution among the clades, a good agreement for species determination was observed among all these genetic classification criteria (see Figs. 1-3). The origin of MP isolates is unclear, although the most plausible explanation is that they may originate from bacteria inadvertently carried by the author owner of this garden. Anyway, these results underscore the high adaptability of *Bradyrhizobium* spp. to soils, both from the point of view of its persistence in soil and genotypic plasticity, as reported previously in Brazilian soils (Loureiro *et al*., 2009). The MP/S-01 strain that we choose for the studies has a DNA fingerprint band pattern very similar to LH/S-14, but they differed in symbiotic behavior. While LH/S-14 was among the worst as revealed by PCA, shoot N accumulation and chlorophyll contents, this isolate was curious in that it produced very few nodules, although with high individual nodule dry weight. By contrast, MP/S-01 had a behavior not so bad, being the 4^th^ among the isolates in shoot N and chlorophyll contents and the 7^th^ in total nodules dry weight (for details, see Supplementary Table S2). In addition, there were 7 single nucleotide differences between the 16S rDNA sequences obtained from LH/S-14 and MP/S-01 PCR products (not shown), whereby the similarity in DNA fingerprint band pattern observed between these isolates seems not to indicate genetic identity.

The genetic diversity observed among our isolates was reflected in disparate symbiotic performances, especially regarding total nodules dry weight and shoot N. The presence of such a diversity in symbiotic performance of local isolates was already described (Damirgi *et al*., 1967, Ellis *et al*., 1984; Keyser and Cregan, 1987; Hirsch, 1996; Rodríguez-Echeverría *et al*., 2003; Abaidoo *et al*., 2007; Melchiorre *et al*., 2011; Iturralde *et al*., 2019), and set the basis for the hypothesis that local strains raise a competitive barrier to nodulation by elite inoculant strains (Dowling and Broughton, 1986; Triplett and Sadowsky, 1992; Pérez-Giménez *et al*., 2011; Checcucci *et al*., 2017; Onishchuk *et al*., 2017; Mendoza-Suárez *et al*., 2021). Moreover, it was repeatedly observed that elite genotypes are absent from local populations, even if the elite strains are continuously used as inoculants, indicating that they possess a poor adaptability to local soil conditions (Loureiro *et al*., 2007; Melchiorre *et al*., 2011; Iturralde *et al*., 2019). This was again observed here, because the genotypic pattern of *B. japonicum* E109, the most used strain in inoculants in Argentina, was not represented among the isolated genotypes.

The diversity of symbiotic performance enabled us to search for correlations between symbiotic parameters and chlorophyll contents. We observed high correlations between shoot N and chlorophyll contents as well as total nodules dry weight and chlorophyll contents, both of which, along with PCA, allowed the identification of *B. diazoefficiens* LH/S-11 and LH/S-12 as the most promising isolates. Since differences in performing the whole denitrification process from oxidized N-compounds to N_2_ are known between the different *Bradyrhizobium* species (Siqueira *et al*., 2017; Obando *et al*., 2019; Mania *et al*., 2020), we looked for the presence of the *nosZ* gene encoding the N_2_O reductase in our isolates as an additional and independent quality criterion. In agreement with previous reports (Siqueira *et al*., 2017; Obando *et al*., 2019), we observed that the three *B. diazoefficiens* isolates possessed at least one copy of *nosZ*, while the *B. elkanii* and *B. japonicum* isolates did not. Therefore, we may expect that inoculation of soybean with *B. diazoefficiens* LH/S- 11 and LH/-12 may prevent N_2_O emissions caused by the rhizobia from this crop, and even reduce such emissions if the inoculation technology fosters high nodules occupation by the inoculated strain in competition with soybean-nodulating strains from the soil microbiota (López-García 2009; Akiyama *et al*., 2016).

The *B. elkanii* isolate LH/S-02 was the only one of this species that possessed a good symbiotic performance, near *B. diazoefficiens* LH/S-11 and LH/S-12. This isolate shared with the other two *B. elkanii* the property of producing and secreting IAA, which was not detected in the other species tested here. In addition, LH/S-02 was the top producer of IAA, although at less than half the level of *A. argentinense* Az39, which is a well-known plant growth promoting rhizobacterium in part due to this capacity (Molina *et al*., 2018). Nevertheless, IAA production and secretion were observed *in vitro*, whereby we cannot extrapolate it to the situation in the rhizosphere or inside plant tissues (e.g., infection threads or nodules), but the possibility exists that the good performance of LH/S-02 be due in part to this capability. On the other hand, none of the isolates produced siderophores or solubilized phosphate at appreciable levels, by difference with other *Bradyrhizobium* spp. field isolates reported elsewhere (Ibny *et al*., 2019; Bünger *et al*., 2021; Bharti *et al*., 2023; Lamrabet *et al*., 2023).

## 5. Conclusion

The symbiotic performance of a soybean crop growing in a soil populated by local, allochtonous genotypes of soybean-nodulating rhizobia would depend on the local strains more than on the elite strains inoculated at sowing. Therefore, it is advisable to use good strains adapted to the local conditions instead of a foreign elite strain. Our results suggest that, to select and assess the quality of such strains, the use of a chlorophyllometer and the PCR-determination of the presence of *nosZ* are advisable methods to incorporate in routine quality control protocols at the inoculants industry. On the one hand, the chlorophyllometer is a small, portable equipment with a moderate cost and no need for consumables, which makes it suitable for small and medium-sized inoculants enterprises, and on the other hand, PCR is already a routine method in most laboratories and inoculants industries. Therefore, selecting strains from local populations and adding these two techniques to the routine quality control analysis of inoculants are three simple measures that may render substantial benefits for profiting N_2_-fixation while, at the same time, mitigating N_2_O greenhouse gas emissions. Undertaking this simple strategy may positively impact on sustainable and climate-friendly soybean agriculture.

## 7. Author contributions

Damián Brignoli: Investigation; methodology; formal analysis; data curation; validation; visualization; writing - review & editing.

Emilia Frickel-Critto: Investigation; methodology; validation; visualization. Tamara J. Sandobal: Investigation; methodology.

Rocío S. Balda: Investigation; methodology.

Cecilia B. Castells: Conceptualization; investigation; methodology; validation; visualization. Elías J. Mongiardini: Conceptualization; investigation; methodology; validation.

Julieta Pérez-Giménez: Conceptualization; investigation; methodology; supervision; validation; visualization; writing - review & editing.

Aníbal R. Lodeiro: Conceptualization; investigation; methodology; funding acquisition; project administration; resources; supervision; validation; visualization; writing - original draft, review & editing.

## 8. Declaration of generative AI and AI-assisted technologies in the writing process

During the preparation of this work the authors used Chat GPT in order to improve the grammar of some paragraphs in the Introduction. After using this tool, the authors reviewed and edited the content as needed and take full responsibility for the content of the publication.

## 9 Acknowledgments

The authors are grateful to Luciana Cayuela and Silvana Stongiani for laboratory technical assistance, Ulises Mancini for help with revision of DNA sequences, and Claudio Mazo for assistance in plant experiments. This work was supported by Agencia Nacional de Promoción de la Investigación, el Desarrollo Tecnológico y la Innovación (Argentina) [grant number PICT2020- 3911]. This founder had no role in study design; in the collection, analysis and interpretation of data; in the writing of the report; and in the decision to submit the article for publication. DB, EFC, TJS, and RSB are fellows of CONICET (Argentina). CBC, EJM, JPG and ARL are members of the Scientific Researcher Career of CONICET (Argentina).

## Abbreviations

ANOVA: analysis of variance
HPLC: high-performance liquid chromatography
IAA: indoleacetic acid
PCA: principal components analysis
SPAD: soil-plant analysis development
UPMGA: unweighted pair-group method with arithmetic mean.

## Supplementary information

**Table S1:** GenBank accession numbers of DNA sequences reported in this work.

**Table S2:** Symbiotic performance of soybean-nodulating isolates.

**Fig. S1:** Genotypic characterization of the soybean-nodulating isolates.

**Fig. S2:** HPLC chromatogram of cellular extracts from the *B. elkanii* isolates.

**Fig. S1.**
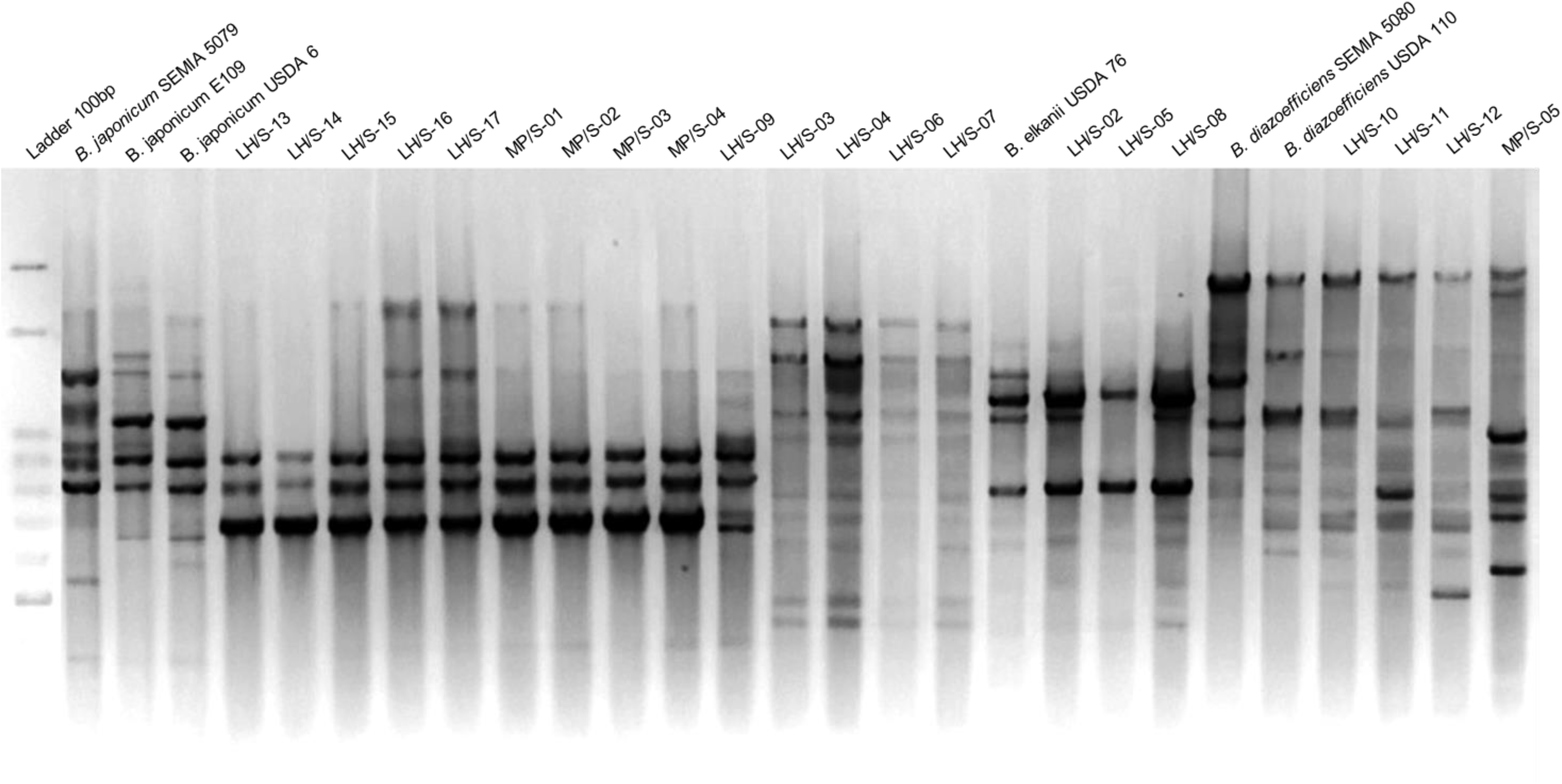
Genotypic characterization of the soybean-nodulating isolates. Total DNA was amplified with the BOX-A1R primer and the banding profile was visualized by agarose 1.5% (w/v) gel electrophoresis revealed with ethidium bromide.

**Fig. S2.**
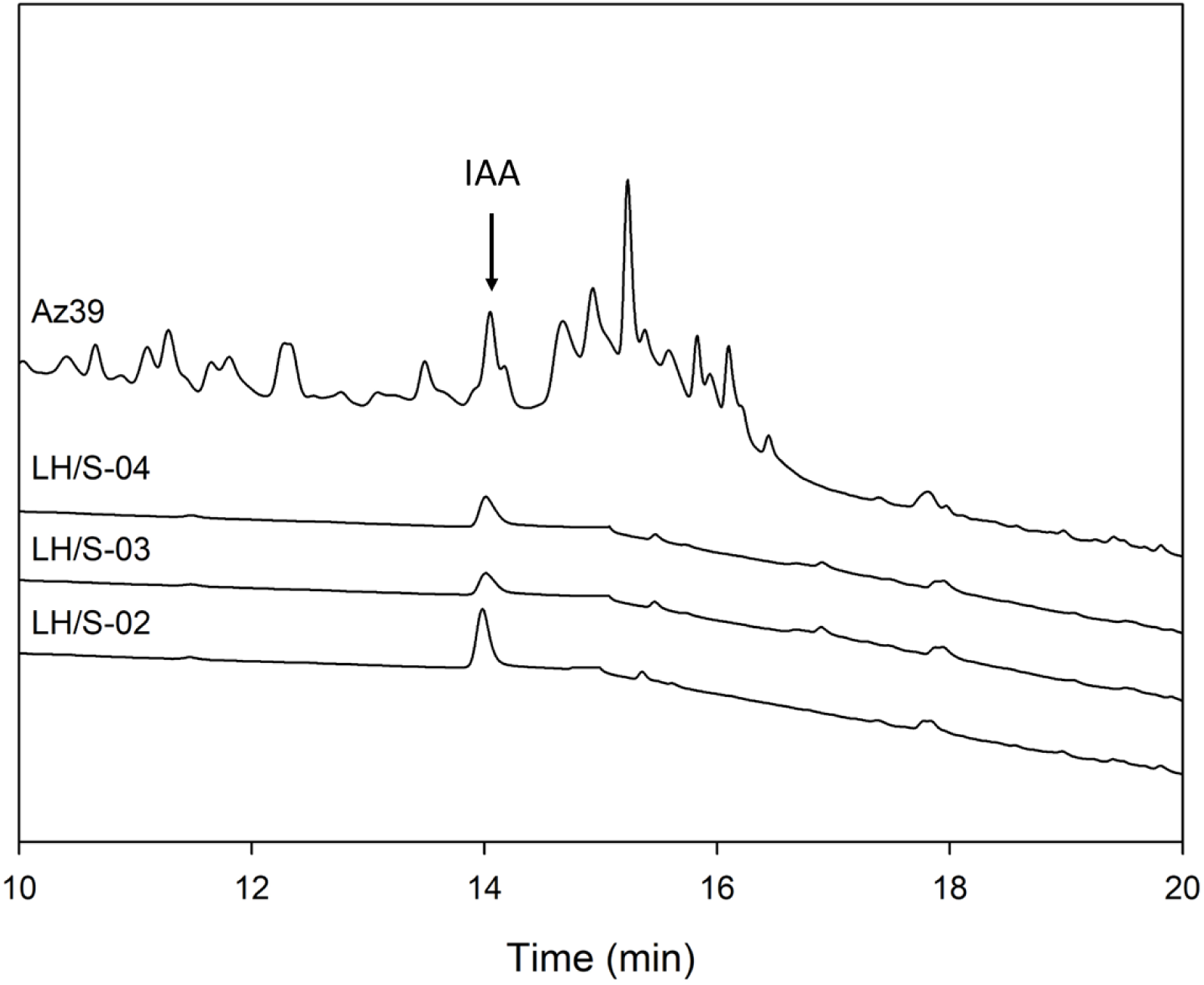
HPLC chromatogram of cellular extracts from *B. elkanii* LH/S-02, LH/S-03 and LH/S-04, and *Azospirillum argentinense* Az39. The arrow indicates the position of a purified indoleacetic acid (IAA) standard.

**Table S1.**
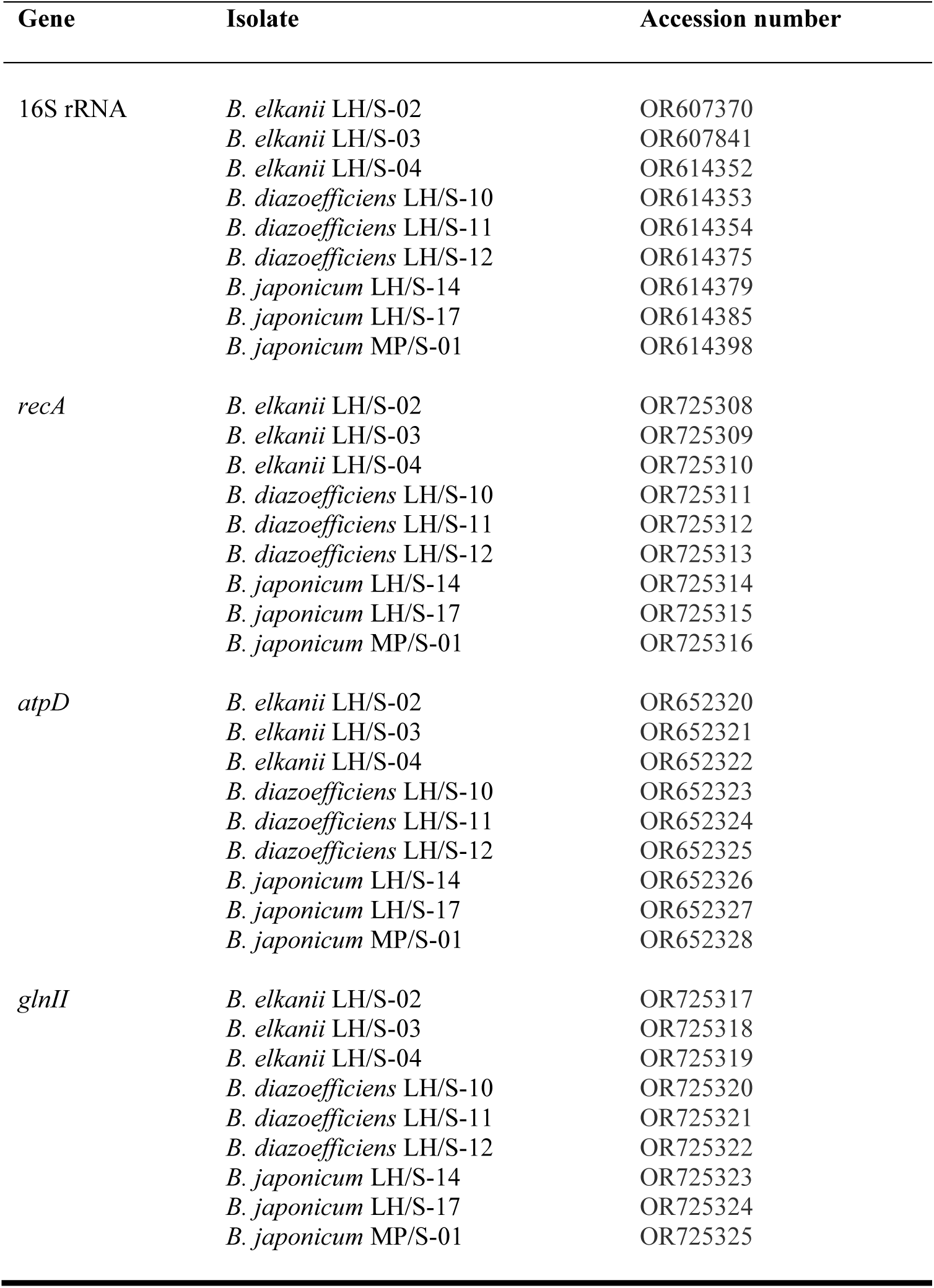
Accession numbers of DNA sequences reported in this work.

| Gene | Isolate | Accession number |
| --- | --- | --- |
| 16S rRNA | *B. elkanii* LH/S-02 | OR607370 |
|  | *B. elkanii* LH/S-03 | OR607841 |
|  | *B. elkanii* LH/S-04 | OR614352 |
|  | *B. diazoefficiens* LH/S-10 | OR614353 |
|  | *B. diazoefficiens* LH/S-11 | OR614354 |
|  | *B. diazoefficiens* LH/S-12 | OR614375 |
|  | *B. japonicum* LH/S-14 | OR614379 |
|  | *B. japonicum* LH/S-17 | OR614385 |
|  | *B. japonicum* MP/S-01 | OR614398 |
| *recA* | *B. elkanii* LH/S-02 | OR725308 |
|  | *B. elkanii* LH/S-03 | OR725309 |
|  | *B. elkanii* LH/S-04 | OR725310 |
|  | *B. diazoefficiens* LH/S-10 | OR725311 |
|  | *B. diazoefficiens* LH/S-11 | OR725312 |
|  | *B. diazoefficiens* LH/S-12 | OR725313 |
|  | *B. japonicum* LH/S-14 | OR725314 |
|  | *B. japonicum* LH/S-17 | OR725315 |
|  | *B. japonicum* MP/S-01 | OR725316 |
| *atpD* | *B. elkanii* LH/S-02 | OR652320 |
|  | *B. elkanii* LH/S-03 | OR652321 |
|  | *B. elkanii* LH/S-04 | OR652322 |
|  | *B. diazoefficiens* LH/S-10 | OR652323 |
|  | *B. diazoefficiens* LH/S-11 | OR652324 |
|  | *B. diazoefficiens* LH/S-12 | OR652325 |
|  | *B. japonicum* LH/S-14 | OR652326 |
|  | *B. japonicum* LH/S-17 | OR652327 |
|  | *B. japonicum* MP/S-01 | OR652328 |
| *glnII* | *B. elkanii* LH/S-02 | OR725317 |
|  | *B. elkanii* LH/S-03 | OR725318 |
|  | *B. elkanii* LH/S-04 | OR725319 |
|  | *B. diazoefficiens* LH/S-10 | OR725320 |
|  | *B. diazoefficiens* LH/S-11 | OR725321 |
|  | *B. diazoefficiens* LH/S-12 | OR725322 |
|  | *B. japonicum* LH/S-14 | OR725323 |
|  | *B. japonicum* LH/S-17 | OR725324 |
|  | *B. japonicum* MP/S-01 | OR725325 |

**Table S2.** Symbiotic performance of soybean-nodulating isolates. *B. japonicum* E109 was included as reference strain.

| Isolate | Nodules<br>plant<sup>-1</sup> | Dry weight (mg) |  |  | Shoot/root<br>ratio (dry) | Fresh weight (mg) |  | Shoot/root<br>ratio (fresh) | Chlorophyll<br>(SPAD units) | Shoot N (mg g<sup>-1</sup><br>dry weight) |
| --- | --- | --- | --- | --- | --- | --- | --- | --- | --- | --- |
|  |  | Individual<br>nodule | Shoot | Root |  | Shoot | Root |  |  |  |
| *B. elkanii* LH/S-02 | 38 d | 1.4 bc | 505.7 abc | 180.0 abc | 2.9 | 2615.7 ab | 1291.4 ab | 2.0 | 30.2 cdef | 28.5 abc |
| *B. elkanii* LH/S-03 | 24 cd | 1.6 bcde | 442.9 abc | 157.1 ab | 3.1 | 2361.4 ab | 1167.1 a | 2.1 | 25.6 abc | 27.6 ab |
| *B. elkanii* LH/S-04 | 24 cd | 1.6 cde | 418.6 ab | 141.4 ab | 3.1 | 2301.4 ab | 1422.9 abc | 1.7 | 27.6 bdc | 37.5 ef |
| *B. diazoefficiens* LH/S-10 | 19 bc | 1.3 abc | 445.7 abc | 141.4 ab | 3.2 | 2511.4 ab | 1468.6 abc | 1.7 | 27.0 abcd | 33.2 bcd |
| *B. diazoefficiens* LH/S-11 | 36 d | 1.2 abc | 530.0 bc | 248.6 d | 2.1 | 2654.3 ab | 1732.9 cde | 1.6 | 33.06 ef | 38.5 f |
| *B. diazoefficiens* LH/S-12 | 28 cd | 1.5 bcd | 571.4 c | 248.6 d | 2.3 | 2734.3 ab | 1767.1 bcde | 1.6 | 34.5 f | 34.1 cde |
| *B. japonicum* LH/S-14 | 8 b | 4.0 de | 360.0 a | 124.3 a | 2.9 | 2184.3 a | 1285.7 ab | 1.7 | 24.4 ab | 28.3 ab |
| *B. japonicum* LH/S-17 | 18 bc | 2.3 de | 517.1 bc | 224.3 cd | 2.3 | 3050.0 b | 1988.6 de | 1.6 | 28.9 bcde | 36.8 def |
| *B. japonicum* MP/S-01 | 15 abc | 2.4 e | 471.4 abc | 177.1 bcd | 2.7 | 2794.3 ab | 1330.0 abc | 2.1 | 29.3 bcde | 37.1 def |
| *B. japonicum* E109 | 24 cd | 1.1 ab | 430.0 abc | 136.0 ab | 3.2 | 2230.0 a | 1468.0 abcd | 1.6 | 30.6 def | 32.8 cd |
| Uninoculated control | 0 | 0 | 498.0 abc | 184.0 bcd | 2.7 | 2200.0 a | 2218.0 e | 1.0 | 22.3 a | 14.5 a |

Plants were grown for 28 days in a plant growth chamber at 30 °C/20 °C day/night temperature, 60%/80% day/night relative humidity, and 16 h photophase. Columns represent means for seven plants (N = 7). Statistical analysis was performed by ANOVA followed by Tukey test. Values followed by the same letter were similar with $\alpha = 0.05$ .

